## Supplementary Materials for "Dynamic expectations: Behavioral and electrophysiological evidence of sub-second updates in reward predictions"

#### Ratings analyses (EEG Studies 1 and 3)

Participants' raw ratings were converted to a standardized z score, based on each participant's mean and standard deviation for that rating. To test whether participants' happiness and motivation ratings varied as a function of their outcome, we ran mixed model linear regressions with a random effect of outcome, encoded as three dummy variables (Win, Near Win After, Near Win Before; Full Miss serving as a baseline), and a fixed effect of participant. To account for the length of the experiment and the fact that participants reported being less engaged as the game progressed, we ran these regressions separately for the first and the second half of the experiment.

**Study 1:** Throughout the experiment, participants were happier and more willing to play again after Wins than after all types of misses (Happiness, first half: Win: 1.01 ± 0.10; NWB: -0.16 ± 0.09; NWA: 0.10 ± 0.11; FM: 0.08 ± 0.06; all p's<.001; second Half: Win: 0.64 ± 0.10; NWB: -0.35 ± 0.09; NWA: -0.37 ± 0.08; FM: -0.32 ± 0.05, all p's<.001; Motivation, first half: Win: 0.90 ± 0.11; NWB: .25 ± 0.10; NWA: 0.44 ± 0.11; FM: 0.32 ± 0.06; all p's<=.001; second half: (Win: 0.02 ± 0.10; NWB: -0.46 ± 0.08; NWA: -0.51 ± 0.08; FM: -0.45 ± 0.05, all p's<.001). In addition, in the first half of the experiment, participants were less happy following NWB compared to NWA (p=0.051) and FM (p=0.025), and more willing to play again after NWA than NWB (p=.096). There were no differences between the different types of misses in the second half (all p's>0.7). These results are in line with previous studies showing that Near Win (Before and After pooled together) are less pleasant than Full Misses but increase willingness to play again[1–4]. The few studies separating NWB from NWA report conflicting effects, with some studies showing that these two effects are mostly driven by Near Win After[2,5], and others showing different effects of NWB and NWA[6]. The fact these Near Wins effects disappeared in the second half of the experiment suggests that they are temporally limited, or that participants got bored or tired of the game at some point.

**Study 2:** The behavioral results were similar to those of Study 1. Throughout the experiment, participants were less happy and less willing to play again after Losses than after all types of Escapes (Happiness, first half: Loss: -0.67 ± 0.11; NLB: 0.68 ± 0.10; NLA: 0.21 ± 0.11; FE: 0.25 ± 0.06; all p's<.001; second Half: Loss: -.98 ± 0.07; NLB: -0.10 ± 0.09; NLA: -0.01 ± 0.08; FE: 0.02 ± 0.05, all p's<.001; Motivation, first half: Loss: -0.10 ± 0.13; NLB: .77 ± 0.11; NLA: 0.39 ± 0.09; FE: 0.41 ± 0.05; all p's<=.001; second half: Loss: -0.76 ± 0.10; NLB: -0.34 ± 0.08; NLA: -0.26 ± 0.09; FE: -0.27 ± 0.05, all p's<.001). In addition, in the first half of the experiment, participants were happier and more willing to play again following NLB compared to NLA and FE (p's<.003). There were no differences in the second half (p's>0.229).

#### Supplementary Tables 1 – 8: Study 1's EEG Repeated-Measure ANOVA results

Tukey pairwise comparisons are presented for Repeated-Measure ANOVAs that reached significance (p<.008).

Table 1. Study 1 - Deceleration [-3000 -2500]

| | | | Prob > F | | | |
|---|---|---|---|---|---|---|
| Source | df | F | Regular | H-F | G-G | Box |
| outcome | 3 | 0.89 | 0.4475 | 0.4369 | 0.4306 | 0.3512 |
| Residual | 105 | | | | | |

Table 2. Study 1 - Deceleration [-2500 -2000]

| | | | Prob > F | | | |
|---|---|---|---|---|---|---|
| Source | df | F | Regular | H-F | G-G | Box |
| outcome | 3 | 1.93 | 0.129 | 0.144 | 0.1477 | 0.1734 |
| Residual | 105 | | | | | |

Table 3. Study 1 - Deceleration [-2000 -1500]

| | | | Prob > F | | | |
|---|---|---|---|---|---|---|
| Source | df | F | Regular | H-F | G-G | Box |
| outcome | 3 | 5.72 | 0.0012 | 0.0023 | 0.003 | 0.0223 |
| Residual | 105 | | | | | |

| | | | Tukey | | Tukey | |
|---|---|---|---|---|---|---|
| | Contrast | Std. err. | t | P>t | [95% conf | interval] |
| NWB vs Win | 2.091598 | 0.735839 | 2.84 | 0.027 | 0.170576 | 4.012619 |
| NWA vs Win | -0.84219 | 0.735839 | -1.14 | 0.663 | -2.76322 | 1.078827 |
| FM vs Win | 0.086329 | 0.735839 | 0.12 | 0.999 | -1.83469 | 2.00735 |
| NWA vs NWB | -2.93379 | 0.735839 | -3.99 | 0.001 | -4.85481 | -1.01277 |
| FM vs NWB | -2.00527 | 0.735839 | -2.73 | 0.037 | -3.92629 | -0.08425 |
| FM vs NWA | 0.928523 | 0.735839 | 1.26 | 0.589 | -0.9925 | 2.849544 |

Table 4 - Deceleration [-1500 -1000]

| | | | Prob>F | | | |
|---|---|---|---|---|---|---|
| Source | df | F | Regular | H-F | G-G | Box |
| outcome | 3 | 14.87 | <.001 | <.001 | <.001 | 0.0005 |
| Residual | 105 | | | | | |

| | | | Tukey | | Tukey | |
|---|---|---|---|---|---|---|
| | Contrast | Std. err. | t | P>t | [95% conf | interval] |
| NWB vs Win | 3.682712 | 0.734276 | 5.02 | <.001 | 1.765773 | 5.599652 |
| NWA vs Win | -0.56037 | 0.734276 | -0.76 | 0.871 | -2.47731 | 1.356574 |
| FM vs Win | 2.406731 | 0.734276 | 3.28 | 0.008 | 0.489791 | 4.32367 |
| NWA vs NWB | -4.24308 | 0.734276 | -5.78 | <.001 | -6.16002 | -2.32614 |
| FM vs NWB | -1.27598 | 0.734276 | -1.74 | 0.31 | -3.19292 | 0.640958 |
| FM vs NWA | 2.967096 | 0.734276 | 4.04 | 0.001 | 1.050156 | 4.884036 |

Table 5. Study 1 - Deceleration [-1000 -500]

| | | | Prob > F | | | |
|---|---|---|---|---|---|---|
| Source | df | F | Regular | H-F | G-G | Box |
| outcome | 3 | 27.5 | <.001 | <.001 | <.001 | <.001 |
| Residual | 105 | | | | | |

| | | | Tukey | | Tukey | |
|---|---|---|---|---|---|---|
| | Contrast | Std. err. | t | P>t | [95% conf | interval] |
| NWB vs Win | 4.74195 | 0.797405 | 5.95 | <.001 | 2.6602 | 6.823699 |
| NWA vs Win | -0.04877 | 0.797405 | -0.06 | 1 | -2.13052 | 2.032978 |
| FM vs Win | 5.408381 | 0.797405 | 6.78 | <.001 | 3.326631 | 7.49013 |
| NWA vs NWB | -4.79072 | 0.797405 | -6.01 | <.001 | -6.87247 | -2.70897 |
| FM vs NWB | 0.666431 | 0.797405 | 0.84 | 0.837 | -1.41532 | 2.748181 |
| FM vs NWA | 5.457152 | 0.797405 | 6.84 | <.001 | 3.375403 | 7.538902 |

Table 6. Study 1 - Deceleration [-500 0]

| | | | Prob > F | | | |
|---|---|---|---|---|---|---|
| Source | df | F | Regular | H-F | G-G | Box |
| outcome | 3 | 13.37 | <.001 | <.001 | <.001 | 0.0008 |
| Residual | 105 | | | | | |

| | | | Tukey | | Tukey | |
|---|---|---|---|---|---|---|
| | Contrast | Std. err. | t | P>t | [95% conf | interval] |
| NWB vs Win | 1.044936 | 0.813267 | 1.28 | 0.575 | -1.07822 | 3.168096 |
| NWA vs Win | 4.467786 | 0.813267 | 5.49 | <.001 | 2.344626 | 6.590945 |
| FM vs Win | 3.605879 | 0.813267 | 4.43 | <.001 | 1.482719 | 5.729039 |
| NWA vs NWB | 3.42285 | 0.813267 | 4.21 | <.001 | 1.29969 | 5.546009 |
| FM vs NWB | 2.560943 | 0.813267 | 3.15 | 0.011 | 0.437783 | 4.684103 |
| FM vs NWA | -0.86191 | 0.813267 | -1.06 | 0.715 | -2.98507 | 1.261253 |

Table 7. Study 1 - FRN

| | | | Prob > F | | | |
|---|---|---|---|---|---|---|
| Source | df | F | Regular | H-F | G-G | Box |
| outcome | 3 | 29.31 | <.001 | <.001 | <.001 | <.001 |
| Residual | 105 | | | | | |

| | | | Tukey | | Tukey | |
|---|---|---|---|---|---|---|
| | Contrast | Std. err. | t | P>t | [95% conf | interval] |
| NWB vs Win | -7.80911 | 0.931541 | -8.38 | <.001 | -10.241 | -5.37718 |
| NWA vs Win | -6.84022 | 0.931541 | -7.34 | <.001 | -9.27215 | -4.40829 |
| FM vs Win | -6.48293 | 0.931541 | -6.96 | <.001 | -8.91487 | -4.051 |
| NWA vs NWB | 0.968893 | 0.931541 | 1.04 | 0.726 | -1.46304 | 3.400824 |
| FM vs NWB | 1.326178 | 0.931541 | 1.42 | 0.488 | -1.10575 | 3.758109 |
| FM vs NWA | 0.357285 | 0.931541 | 0.38 | 0.981 | -2.07465 | 2.789216 |

Table 8. Study 1 - P3

| | | | Prob > F | | | |
|---|---|---|---|---|---|---|
| Source | df | F | Regular | H-F | G-G | Box |
| outcome | 3 | 42.57 | <.001 | <.001 | <.001 | <.001 |
| Residual | 105 | | | | | |

| | | | Tukey | | Tukey | |
|---|---|---|---|---|---|---|
| | Contrast | Std. err. | t | P>t | [95% conf | interval] |
| NWB vs Win | -4.51423 | 0.867508 | -5.2 | <.001 | -6.779 | -2.24947 |
| NWA vs Win | -8.83128 | 0.867508 | -10.18 | <.001 | -11.096 | -6.56652 |
| FM vs Win | -7.92357 | 0.867508 | -9.13 | <.001 | -10.1883 | -5.65881 |
| NWA vs NWB | -4.31705 | 0.867508 | -4.98 | <.001 | -6.58181 | -2.05229 |
| FM vs NWB | -3.40934 | 0.867508 | -3.93 | 0.001 | -5.67411 | -1.14458 |
| FM vs NWA | 0.907706 | 0.867508 | 1.05 | 0.723 | -1.35706 | 3.17247 |

### Supplementary Figure 1 – Study 1's FRN and P3

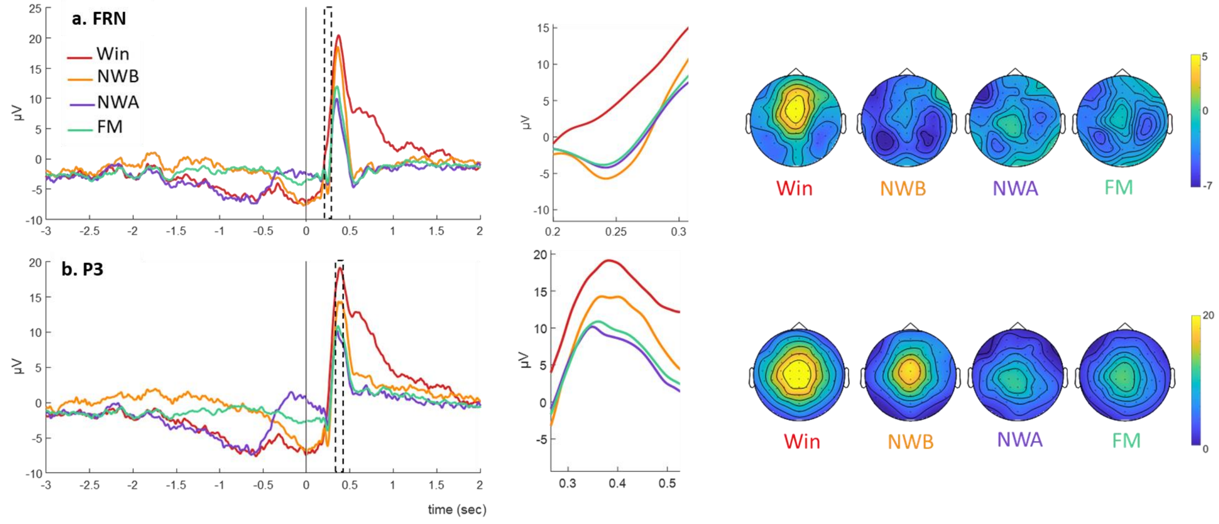

**Supplementary Figure 1: Study 1's EEG results. a. FRN results:** Left panel shows the grand average ERPs for each outcome locked to the stop of the machine at FRN electrodes of interest pooled (Fz, FCz, Cz). The dashed rectangle over the waveforms indicates the time window used for the FRN analysis. Middle panel shows a zoom in of that time window. Right panel shows the topographies for each outcome. **b. P3 results:** Same for P3, for average of electrodes Cz, CPz, Pz, POz, Oz.

### Supplementary Tables 8-14: Study 3's EEG Repeated-Measure ANOVA results

Table 8. Study 3 - Deceleration [-3000 -2500]

| | | | Prob > F | | | |
|---|---|---|---|---|---|---|
| Source | df | F | Regular | H-F | G-G | Box |
| outcome | 3 | 1.43 | 0.2371 | 0.2397 | 0.2417 | 0.2393 |
| Residual | 102 | | | | | |

Table 9. Study 3 - Deceleration [-2500 -2000]

| | | | Prob > F | | | |
|---|---|---|---|---|---|---|
| Source | df | F | Regular | H-F | G-G | Box |
| outcome | 3 | 3.53 | 0.0175 | 0.0246 | 0.0279 | 0.0688 |
| Residual | 102 | | | | | |

Table 10. Study 3 - Deceleration [-2000 -1500]

| | | | Prob > F | | | |
|---|---|---|---|---|---|---|
| Source | df | F | Regular | H-F | G-G | Box |
| outcome | 3 | 7.48 | <.001 | <.001 | <.001 | 0.0098 |
| Residual | 102 | | | | | |

| | | | Tukey | | Tukey | |
|---|---|---|---|---|---|---|
| | Contrast | Std. err. | t | P>t | [95% conf | interval] |
| NLB vs Loss | 0.94813 | 0.720067 | 1.32 | 0.554 | -0.93261 | 2.828867 |
| NLA vs Loss | -1.61891 | 0.720067 | -2.25 | 0.117 | -3.49965 | 0.261823 |
| FE vs Loss | -2.03004 | 0.720067 | -2.82 | 0.029 | -3.91077 | -0.1493 |
| NLA vs NLB | -2.56704 | 0.720067 | -3.57 | 0.003 | -4.44778 | -0.68631 |
| FE vs NLB | -2.97817 | 0.720067 | -4.14 | <.001 | -4.8589 | -1.09743 |
| FE vs NLA | -0.41112 | 0.720067 | -0.57 | 0.941 | -2.29186 | 1.469614 |

Table 11. Study 3 - Deceleration [-1500 -1000]

| | | | Prob>F | | | |
|---|---|---|---|---|---|---|
| Source | df | F | Regular | H-F | G-G | Box |
| outcome | 3 | 9.31 | <.001 | <.001 | 0.0001 | 0.0044 |
| Residual | 102 | | | | | |

| | | | Tukey | | Tukey | |
|---|---|---|---|---|---|---|
| | Contrast | Std. err. | t | P>t | [95% conf | interval] |
| NLB vs Loss | 1.305904 | 0.707803 | 1.85 | 0.259 | -0.5428 | 3.154611 |
| NLA vs Loss | -2.31367 | 0.707803 | -3.27 | 0.008 | -4.16237 | -0.46496 |
| FE vs Loss | -0.94363 | 0.707803 | -1.33 | 0.544 | -2.79234 | 0.905077 |
| NLA vs NLB | -3.61957 | 0.707803 | -5.11 | <.001 | -5.46828 | -1.77087 |
| FE vs NLB | -2.24953 | 0.707803 | -3.18 | 0.01 | -4.09824 | -0.40083 |
| FE vs NLA | 1.370038 | 0.707803 | 1.94 | 0.22 | -0.47867 | 3.218745 |

Table 12. Study 3 - Deceleration [-1000 -500]

| | | | Prob > F | | | |
|---|---|---|---|---|---|---|
| Source | df | F | Regular | H-F | G-G | Box |
| outcome | 3 | 17.08 | <.001 | <.001 | <.001 | 0.0002 |
| Residual | 102 | | | | | |

| | | | Tukey | | Tukey | |
|---|---|---|---|---|---|---|
| | Contrast | Std. err. | t | P>t | [95% conf | interval] |
| NLB vs Loss | 2.596125 | 0.807639 | 3.21 | 0.009 | 0.486659 | 4.70559 |
| NLA vs Loss | -2.44756 | 0.807639 | -3.03 | 0.016 | -4.55703 | -0.3381 |
| FE vs Loss | 2.356701 | 0.807639 | 2.92 | 0.022 | 0.247235 | 4.466166 |
| NLA vs NLB | -5.04369 | 0.807639 | -6.24 | <.001 | -7.15315 | -2.93422 |
| FE vs NLB | -0.23942 | 0.807639 | -0.3 | 0.991 | -2.34889 | 1.870041 |
| FE vs NLA | 4.804262 | 0.807639 | 5.95 | <.001 | 2.694796 | 6.913727 |

Table 13. Study 3 - Deceleration [-500 0]

| | | | Prob > F | | | |
|---|---|---|---|---|---|---|
| Source | df | F | Regular | H-F | G-G | Box |
| outcome | 3 | 8.83 | <.001 | <.001 | 0.0001 | 0.0054 |
| Residual | 102 | | | | | |

| | | | Tukey | | Tukey | |
|---|---|---|---|---|---|---|
| | Contrast | Std. err. | t | P>t | [95% conf | interval] |
| NLB vs Loss | 0.198744 | 0.829179 | 0.24 | 0.995 | -1.96698 | 2.364472 |
| NLA vs Loss | 3.32327 | 0.829179 | 4.01 | 0.001 | 1.157542 | 5.488998 |
| FE vs Loss | 2.870044 | 0.829179 | 3.46 | 0.004 | 0.704316 | 5.035772 |
| NLA vs NLB | 3.124526 | 0.829179 | 3.77 | 0.002 | 0.958798 | 5.290255 |
| FE vs NLB | 2.671301 | 0.829179 | 3.22 | 0.009 | 0.505572 | 4.837029 |
| FE vs NLA | -0.45323 | 0.829179 | -0.55 | 0.947 | -2.61895 | 1.712502 |

Table 14. Study 3 - FRN

| | | | Prob > F | | | |
|---|---|---|---|---|---|---|
| Source | df | F | Regular | H-F | G-G | Box |
| outcome | 3 | 26.22 | <.001 | <.001 | <.001 | <.001 |
| Residual | 102 | | | | | |

| | | | Tukey | | Tukey | |
|---|---|---|---|---|---|---|
| | Contrast | Std. err. | t | P>t | [95% conf | interval] |
| NLB vs Loss | 5.869042 | 0.952402 | 6.16 | <.001 | 3.381469 | 8.356615 |
| NLA vs Loss | 7.712022 | 0.952402 | 8.1 | <.001 | 5.224449 | 10.19959 |
| FE vs Loss | 6.604507 | 0.952402 | 6.93 | <.001 | 4.116934 | 9.09208 |
| NLA vs NLB | 1.84298 | 0.952402 | 1.94 | 0.22 | -0.64459 | 4.330553 |
| FE vs NLB | 0.735465 | 0.952402 | 0.77 | 0.867 | -1.75211 | 3.223038 |
| FE vs NLA | -1.10752 | 0.952402 | -1.16 | 0.652 | -3.59509 | 1.380058 |

Table 14. Study 3 - P3

| | | | Prob > F | | | |
|---|---|---|---|---|---|---|
| Source | df | F | Regular | H-F | G-G | Box |
| outcome | 3 | 31.52 | <.001 | <.001 | <.001 | <.001 |
| Residual | 102 | | | | | |

| | | | Tukey | | Tukey | |
|---|---|---|---|---|---|---|
| | Contrast | Std. err. | t | P>t | [95% conf | interval] |
| NLB vs Loss | -1.49612 | 1.053267 | -1.42 | 0.49 | -4.24714 | 1.254904 |
| NLA vs Loss | -8.29418 | 1.053267 | -7.87 | <.001 | -11.0452 | -5.54316 |
| FE vs Loss | -7.48593 | 1.053267 | -7.11 | <.001 | -10.237 | -4.73491 |
| NLA vs NLB | -6.79807 | 1.053267 | -6.45 | <.001 | -9.54909 | -4.04705 |
| FE vs NLB | -5.98982 | 1.053267 | -5.69 | <.001 | -8.74084 | -3.2388 |
| FE vs NLA | 0.80825 | 1.053267 | 0.77 | 0.869 | -1.94277 | 3.55927 |

**Supplementary Fig. 2 – Study 3's FRN and P3**

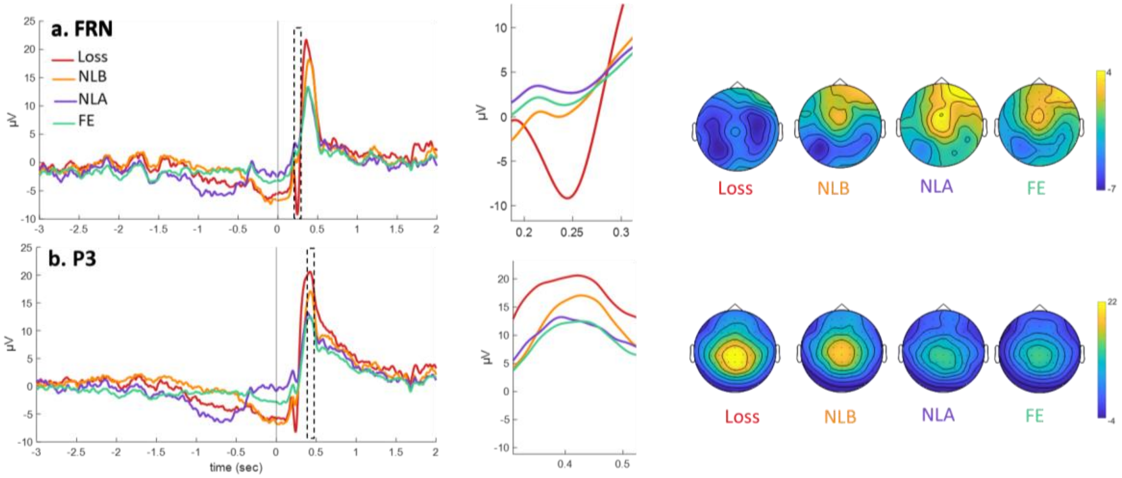

**Supplementary Figure 2: Study 1's EEG results. a. FRN results:** Left panel shows the grand average ERPs for each outcome locked to the stop of the machine at FRN electrodes of interest pooled (Fz, FCz, Cz). The dashed rectangle over the waveforms indicates the time window used for the FRN analysis. Middle panel shows a zoom in of that time window. Right panel shows the topographies for each outcome. **b. P3 results:** Same for P3, for average of electrodes Cz, CPz, Pz, POz, Oz.

### Discussion of FRN and P3 results

In both EEG studies, the Feedback Related Negativity encoded valence with enhanced negative amplitudes for Misses/Losses vs. Wins/Escapes, confirming that participants understood the meaning of a match in both versions of the games. This is consistent with past studies showing that the FRN differentiates between gains and losses[7,8]. However, the FRN did not differentiate between Near Win/Near Loss and Full Miss/Full Escape. The EEG literature on Near Win is divided on this point, with studies finding as in our studies similar FRN for Near Wins and Full Misses[4], and other reporting smaller (less negative) FRN for Near Wins vs. Full Misses[9–11]. These discrepancies may be due to different baseline correction procedures: baseline correcting to the pre-outcome phase might inject expectation differences into the FRN. The lack of FRN sensitivity to the different types of Near Outcomes also contrasts with past research showing the FRN's sensitivity to expectations[12–16]. In these studies, expectations were elicited ahead of the feedback eliciting the FRN, for example with a single cue given a few seconds before outcome onset. Dynamic expectations change up to outcome onset, and the integration of outcome value with expectations may occur later than in the FRN time window. Indeed, when feedback is complex, processes associated with the FRN can be delayed [17,18].

P3 has been associated with surprise and reward prediction errors[19–21]. In Study 1, P3 was larger for NWB compared to NWA and FM[22], suggesting that participants were more surprised by their loss following NWB. This result is in line with the claim that NWB create higher expectations than NWA and FM. Study 3's results were strikingly similar, with larger P3 for NLB vs. NLA and FE. The combined results of Studies 1 and 3 support the finding that P3 tracks unsigned (non-valenced) reward prediction error magnitude[20,23,24]. In addition, P3 was found to be larger for Wins vs. all Misses, consistent with past Near Win studies[4,10,22,25,26]. By themselves, these results suggest that the P3 is encoding reward valence in addition to RPE, with gains eliciting larger P3 than losses [12,16,21,27]. However, in Study 3, P3 was larger for Losses vs. all escapes, suggesting that valence is not what drives the effect in this task. Rather, we suggest that the difference between Wins and Misses/Loss and Escapes reflect the rarity of matches in the game (16.6 %). Alternatively, this difference could reflect reward magnitude: in both studies, the absolute reward associated with a match ($0.25 in Study 1, -$0.25 in Study 2) was higher than the one associated with a mismatch ($0 in Study 1, $0.10 in Study 2), and past studies have suggested that P3 reflects reward magnitude [24,28,29]. While these two explanations are not mutually exclusive, the deceleration findings as well as the larger P3 elicited by NWB/NLB vs. other misses/escapes support the unsigned RPE hypothesis.

**Supplementary Tables 15– 20: Study 2 "Slot or Not"'s behavioral Repeated-Measure ANOVA results**

Table 15. Study 2 - Deceleration [-3000 -2500]

| | | | Prob > F | | | |
|---|---|---|---|---|---|---|
| Source | df | F | Regular | H-F | G-G | Box |
| outcome | 3 | 0.21 | 0.8885 | 0.8696 | 0.8518 | 0.6493 |
| Residual | 87 | | | | | |

Table 16. Study 2 - Deceleration [-2500 -2000]

| | | | Prob > F | | | |
|---|---|---|---|---|---|---|
| Source | df | F | Regular | H-F | G-G | Box |
| outcome | 3 | 0.49 | 0.6874 | 0.6574 | 0.641 | 0.4878 |
| Residual | 87 | | | | | |

Table 17. Study 2 - Deceleration [-2000 -1500]

| | | | Prob > F | | | |
|---|---|---|---|---|---|---|
| Source | df | F | Regular | H-F | G-G | Box |
| outcome | 3 | 1.61 | 0.1923 | 0.1967 | 0.2008 | 0.2142 |
| Residual | 87 | | | | | |

Table 18. Study 2 - Deceleration [-1500 -1000]

| | | | Prob > F | | | |
|---|---|---|---|---|---|---|
| Source | df | F | Regular | H-F | G-G | Box |
| outcome | 3 | 11.34 | <.001 | <.001 | <.001 | 0.0022 |
| Residual | 87 | | | | | |

| | | | Tukey | | Tukey | |
|---|---|---|---|---|---|---|
| | Contrast | Std. err. | t | P>t | [95% conf | interval] |
| NWB vs Win | -0.06833 | 0.03848 | -1.78 | 0.292 | -0.16913 | 0.032461 |
| NWA vs Win | 0.144444 | 0.03848 | 3.75 | 0.002 | 0.04365 | 0.245239 |
| FM vs Win | -0.02056 | 0.03848 | -0.53 | 0.95 | -0.12135 | 0.080239 |
| NWA vs NWB | 0.212778 | 0.03848 | 5.53 | <.001 | 0.111984 | 0.313572 |
| FM vs NWB | 0.047778 | 0.03848 | 1.24 | 0.602 | -0.05302 | 0.148572 |
| FM vs NWA | -0.165 | 0.03848 | -4.29 | <.001 | -0.26579 | -0.06421 |

Table 19. Study 2 - Deceleration [-1000 -500]

| | | | Prob > F | | | |
|---|---|---|---|---|---|---|
| Source | df | F | Regular | H-F | G-G | Box |
| outcome | 3 | 33.63 | <.001 | <.001 | <.001 | <.001 |
| Residual | 87 | | | | | |

| | | | Tukey | | Tukey | |
|---|---|---|---|---|---|---|
| | Contrast | Std. err. | t | P>t | [95% conf | interval] |
| NWB vs Win | -0.17056 | 0.033 | -5.17 | <.001 | -0.257 | -0.08412 |
| NWA vs Win | 0.096111 | 0.033 | 2.91 | 0.023 | 0.009671 | 0.182552 |
| FM vs Win | -0.18167 | 0.033 | -5.51 | <.001 | -0.26811 | -0.09523 |
| NWA vs NWB | 0.266667 | 0.033 | 8.08 | <.001 | 0.180226 | 0.353107 |
| FM vs NWB | -0.01111 | 0.033 | -0.34 | 0.987 | -0.09755 | 0.075329 |
| FM vs NWA | -0.27778 | 0.033 | -8.42 | <.001 | -0.36422 | -0.19134 |

Table 20. Study 2 - Deceleration [-500 0]

| | | | Prob > F | | | |
|---|---|---|---|---|---|---|
| Source | df | F | Regular | H-F | G-G | Box |
| outcome | 3 | 19.06 | <.001 | <.001 | <.001 | 0.0001 |
| Residual | 87 | | | | | |

| | | | Tukey | | Tukey | |
|---|---|---|---|---|---|---|
| | Contrast | Std. err. | t | P>t | [95% conf | interval] |
| 2 vs 1 | -0.23278 | 0.048683 | -4.78 | <.001 | -0.3603 | -0.10526 |
| 3 vs 1 | -0.205 | 0.048683 | -4.21 | <.001 | -0.33252 | -0.07748 |
| 4 vs 1 | -0.36333 | 0.048683 | -7.46 | <.001 | -0.49085 | -0.23581 |
| 3 vs 2 | 0.027778 | 0.048683 | 0.57 | 0.941 | -0.09974 | 0.155299 |
| 4 vs 2 | -0.13056 | 0.048683 | -2.68 | 0.043 | -0.25808 | -0.00303 |
| 4 vs 3 | -0.15833 | 0.048683 | -3.25 | 0.009 | -0.28585 | -0.03081 |

**Supplementary Tables 21– 26: Study 4 "Slot or Not"'s behavioral Repeated-Measure ANOVA results**

Table 21. Study 4 - Deceleration [-3000 -2500]

| | | | Prob > F | | | |
|---|---|---|---|---|---|---|
| Source | df | F | Regular | H-F | G-G | Box |
| outcome | 3 | 0.02 | 0.9972 | 0.9954 | 0.9917 | 0.9006 |
| Residual | 60 | | | | | |

Table 22 .Study 4 - Deceleration [-2500 -2000]

| | | | Prob > F | | | |
|---|---|---|---|---|---|---|
| Source | df | F | Regular | H-F | G-G | Box |
| outcome | 3 | 0.14 | 0.9363 | 0.8751 | 0.8568 | 0.7131 |
| Residual | 60 | | | | | |

Table 23. Study 4 - Deceleration [-2000 -1500]

| | | | Prob > F | | | |
|---|---|---|---|---|---|---|
| Source | df | F | Regular | H-F | G-G | Box |
| outcome | 3 | 1.21 | 0.3126 | 0.3089 | 0.3069 | 0.2837 |
| Residual | 60 | | | | | |

Table 24. Study 4 - Deceleration [-1500 -1000]

| | | | Prob > F | | | |
|---|---|---|---|---|---|---|
| Source | df | F | Regular | H-F | G-G | Box |
| outcome | 3 | 6.68 | 0.0006 | 0.0015 | 0.0023 | 0.0177 |
| Residual | 60 | | | | | |

| | | | Tukey | | Tukey | |
|---|---|---|---|---|---|---|
| | Contrast | Std. err. | t | P>t | [95% conf | interval] |
| NLB vs Loss | -0.00476 | 0.061671 | -0.08 | 1 | -0.16773 | 0.158206 |
| NLA vs Loss | -0.21429 | 0.061671 | -3.47 | 0.005 | -0.37725 | -0.05132 |
| FE vs Loss | 0.031217 | 0.061671 | 0.51 | 0.957 | -0.13175 | 0.194185 |
| NLA vs NLB | -0.20952 | 0.061671 | -3.4 | 0.006 | -0.37249 | -0.04656 |
| FE vs NLB | 0.035979 | 0.061671 | 0.58 | 0.937 | -0.12699 | 0.198946 |
| FE vs NLA | 0.245503 | 0.061671 | 3.98 | 0.001 | 0.082535 | 0.40847 |

Table 25. Study 4 - Deceleration [-1000 -500]

| | | | Prob > F | | | |
|---|---|---|---|---|---|---|
| Source | df | F | Regular | H-F | G-G | Box |
| outcome | 3 | 10.99 | 0 | 0 | 0.0001 | 0.0035 |
| Residual | 60 | | | | | |

| | | | Tukey | | Tukey | |
|---|---|---|---|---|---|---|
| | Contrast | Std. err. | t | P>t | [95% conf | interval] |
| NLB vs Loss | 0.144444 | 0.067318 | 2.15 | 0.151 | -0.03344 | 0.322333 |
| NLA vs Loss | -0.10794 | 0.067318 | -1.6 | 0.385 | -0.28582 | 0.069952 |
| FE vs Loss | 0.250529 | 0.067318 | 3.72 | 0.002 | 0.072641 | 0.428417 |
| NLA vs NLB | -0.25238 | 0.067318 | -3.75 | 0.002 | -0.43027 | -0.07449 |
| FE vs NLB | 0.106085 | 0.067318 | 1.58 | 0.4 | -0.0718 | 0.283973 |
| FE vs NLA | 0.358466 | 0.067318 | 5.32 | <.001 | 0.180577 | 0.536354 |

Table 26. Study 4 - Deceleration [-500 0]

| | | | Prob > F | | | |
|---|---|---|---|---|---|---|
| Source | df | F | Regular | H-F | G-G | Box |
| outcome | 3 | 11.23 | 0 | 0 | 0.0001 | 0.0032 |
| Residual | 60 | | | | | |

| | | | Tukey | | Tukey | |
|---|---|---|---|---|---|---|
| | Contrast | Std. err. | t | P>t | [95% conf | interval] |
| NLB vs Loss | 0.206349 | 0.071026 | 2.91 | 0.026 | 0.018662 | 0.394036 |
| NLA vs Loss | 0.184127 | 0.071026 | 2.59 | 0.056 | -0.00356 | 0.371814 |
| FE vs Loss | 0.411376 | 0.071026 | 5.79 | <.001 | 0.223689 | 0.599063 |
| NLA vs NLB | -0.02222 | 0.071026 | -0.31 | 0.989 | -0.20991 | 0.165465 |
| FE vs NLB | 0.205026 | 0.071026 | 2.89 | 0.027 | 0.01734 | 0.392713 |
| FE vs NLA | 0.227249 | 0.071026 | 3.2 | 0.012 | 0.039562 | 0.414936 |
